# Ultrasound Tracking Reveals Progressive Regional Strain Differences in Human Achilles Tendons During Fatigue Loading

**DOI:** 10.64898/2026.09.09.750111

**Authors:** Sittinon Nuethong, Josh R. Baxter

## Abstract

Ultrasound is commonly used to assess structural changes in symptomatic Achilles tendons, but quantitative biomechanical metrics for progressive tendon deterioration remain limited. The goal of this study was to develop and validate an automated ultrasound tracking algorithm for regional tendon deformation and evaluate strain progression in survived and ruptured tendons during fatigue loading. We hypothesized that maximum strain, average strain, and strain heterogeneity would exhibit different trajectories between groups. Ten cadaveric Achilles tendons underwent cyclic loading with stress tests every 500 cycles until rupture or 150,000 cycles. Ultrasound images acquired during stress tests were analyzed using an automated tracking algorithm to generate spatially resolved regional strain fields. Ultrasound-derived bulk strain was highly correlated with actuator-derived strain in survived (R^2^ = 0.968 ± 0.017) and ruptured tendons (R^2^ = 0.972 ± 0.014). Maximum and average longitudinal strains progressively diverged between groups across fatigue life (Group x FatigueLife: p = 0.003 and p < 0.0001, respectively). During the first 10,000 cycles, average strain decreased in survived tendons (β = −0.0268%, p = 0.0215) but not ruptured tendons (β = 0.0147%, p = 0.1197), with a significant Group x Cycle interaction (p = 0.0061). This study demonstrates that the algorithm quantified Achilles tendon deformation with high fidelity and enabled spatially resolved strain assessment throughout fatigue loading. Maximum and average strain followed different trajectories between groups, whereas strain heterogeneity did not. Early differences in tendon biomechanics suggest that regional strain behavior may change before pronounced differences in absolute magnitude develop.

## 1. Introduction

Achilles tendon ruptures are one of the most common and debilitating injuries of the lower extremity, often resulting in persistent functional deficits and structural alterations of muscle-tendon unit that require prolonged rehabilitation to restore function.^24,26,31,36^ Even though Achilles tendon ruptures are frequently associated with the underlying tendinopathy attributed to overuse and repetitive mechanical loading, characterizing the progression of the fatigue-induced tendon damage before rupture remains a major clinical challenge.^6^ Current clinical assessments primarily detect structural abnormalities after the symptoms develop and provide limited information regarding progressive mechanical deterioration of the tendon tissue.^14,32^ Consequently, quantitative biomechanical metrics capable of characterizing progressive tissue damage are needed to improve understanding of tendon fatigue and help identify compromised tissues.

Repetitive mechanical loading inflicts progressive fatigue damage within tendon tissue before macroscopic rupture occurs.^19,20,37^ Consequently, the fatigue-induced microstructural damage including collagen fibril degradation and localized fiber rupture alters mechanical response of tendon to loading.^19,20,37,39^ Because tendon deformation reflects the biomechanical response of the tendons to applied load, regional strain, a quantitative biomechanical metric of tissue deformation, has emerged as a promising biomarker for assessing progressive tendon damage. Ultrasound imaging has emerged as a noninvasive modality for characterizing tendon structure and tracking tendon motion in vivo.^1,3,17,28,35^ Recent studies have demonstrated that ultrasound speckle tracking can quantitatively measure tendon displacement to evaluate muscle-tendon elongation, tendon strain, subtendon sliding, and stiffness.^4,5,7,8,12,13^ However, the use of ultrasound-based tracking to quantify regional tendon strain throughout progressive fatigue loading and characterize mechanical changes preceding Achilles tendon rupture remains limited. Because fatigue-induced collagen matrix disruption is expected to reduce local load-bearing capacity, regional deformation promisingly serves as a key biomechanical marker of assessing progressive damage. Therefore, a robust and validated ultrasound-based approach capable of quantifying regional tendon deformation throughout fatigue loading is needed to characterize the progression of tendon damage.

The purpose of this study was to develop and validate a robust automated ultrasound tracking algorithm for quantifying regional tendon deformation during prolonged cyclic loading and stress testing. Using this framework, regional strain progression was compared between survived and ruptured tendons throughout fatigue loading. We hypothesized that the proposed tracking algorithm would demonstrate high tracking fidelity, with ultrasound-derived bulk tendon strains exhibiting strong agreement with actuator-derived bulk tendon strains throughout fatigue life. In addition, we expected ultrasound-derived bulk strain measurements at the final fatigue stage to reproduce the previously reported differences between survived and ruptured tendons measured using actuator-derived strain. We further hypothesized that the maximum, average, and standard deviation of regional longitudinal strain would progressively increase in ruptured tendons compared with survived tendons as fatigue loading progressed.

## 2. Materials & Methods

This study used ultrasound images of the cadaveric tendons from our previous work to validate our automated tracking algorithm and quantify regional strain during fatigue loading. The cadaveric tendon preparation, experimental setup, and mechanical testing protocols have been described in this previous work.^33^ Briefly, ten cadaveric tendons from seven donors (4M, 3F; Age: 60 ± 15) were dissected, sectioned into 30-mm long dog-bone shaped specimens, and clamped in a custom-built tank and pulley system with an ultrasound probe mounted to image the tendon midsubstance. The specimens were loaded up to 150,000 cycles or until rupture using a sinusoidal waveform between 10 and 20 MPa at 1 Hz. After every 500 cycles, continuous B-mode ultrasound images of the tendon midsubstance were acquired at 41 Hz during 2 loading cycles at 0.25 Hz to minimize motion artifact during image acquisition.

After the experiment, a custom script (MATLAB R2025b.) was implemented to track the deformation of the tendon midsubstance from the acquired ultrasound images. A rectangular region of interest (ROI) was manually defined over the midsubstance, where a 12 × 3 grid of kernels was placed inside to define the subregions of the midsubstance for tracking (Fig 1A). Similar to our previous work^16^, each kernel was seeded with point trackers generated by Kanade-Lucas-Tomasi (KLT) point tracking algorithm, which detects trackable image features based on the eigenvalues of the image gradient matrix.^25,34,38^ (Fig 1B). The initial center of each kernel was calculated from the four corner points of the original kernel position.

**Fig 1.**
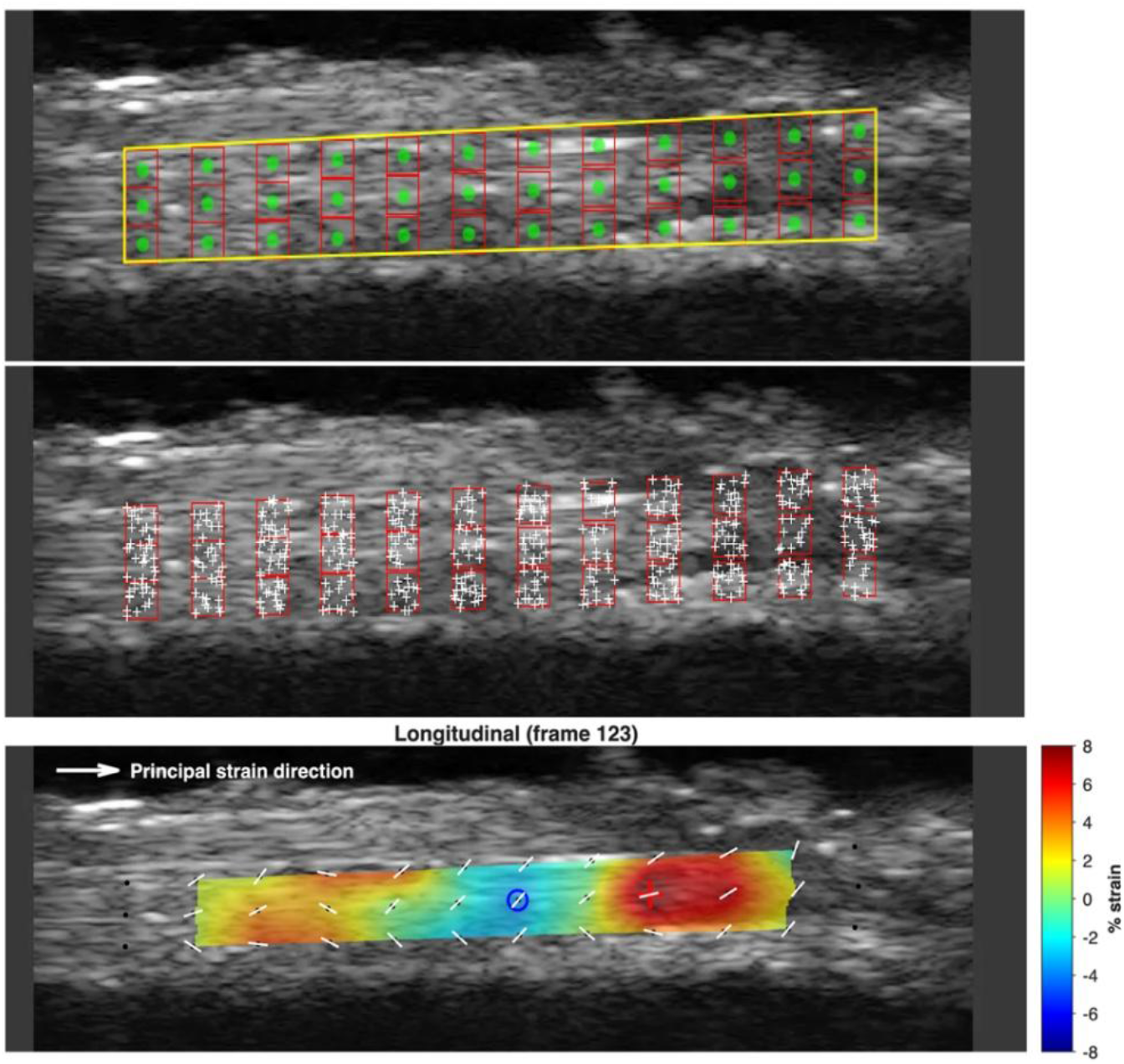
Tracking framework for ultrasound imaging of the cadaveric Achilles tendon. (A) Representative ultrasound image with an ROI (yellow rectangle), a 12 × 3 kernel grid (red rectangles), and kernel center (green dots) within each kernel. (B) Point trackers (white dots) generated by Kanade-Lucas-Tomasi algorithm within each the kernel. (C) Longitudinal strain map of the representative cadaveric Achilles tendon at the peak loading frame. Colors represent local longitudinal strain magnitude (%), and white vectors indicate principal strain directions.

Point trackers in each kernel in each frame returned their updated x- and y-coordinates in the ultrasound image. Matched point-tracker coordinates from the previous and current frames were used to estimate a two-dimensional similarity transformation, which allowed translation, rotation, and uniform scaling of the tracked kernel. This transformation was then applied to the previous kernel center to calculate the new kernel center position in the current frame. Raw frame-to-frame kernel displacement was calculated as the difference in kernel center coordinates between consecutive frames. Due to tendon rotation during loading, principal component analysis (PCA) was used to define local tendon axis and resolve kernel displacements into longitudinal and transverse components^21^. A temporal median filter was applied to reduce frame-to-frame tracking noise, and a spatial moving median filter was applied across neighboring kernels to smooth cumulative displacement fields.

For bulk tendon strain estimation, cumulative longitudinal kernel displacements were first averaged across the tendon depth for each column at every frame. The resulting depth-averaged column displacements were then plotted against the corresponding initial column positions along the tendon axis, and a linear regression was performed. The slope of the regression line was defined as the ultrasound-derived bulk tendon strain, whereas the intercept represented rigid-body translation. Bulk strain values obtained at eleven fatigue stages spanning approximately 1% to 100% fatigue life were compared with actuator-derived bulk tendon strains during the loading phase to evaluate tracking fidelity. To quantify agreement between the two measurements, the toe region of the ultrasound-derived versus actuator-derived strain relationship was excluded using an empirically selected strain cutoff ranging from approximately 0.2% to 1.0%, chosen from the ultrasound-derived versus crosshead-derived strain curves for each specimen. Cutoff selection was applied consistently across fatigue stages within each specimen. The remaining linear region was used to calculate the regression slope, coefficient of determination (R^2^), and root-mean-square error (RMSE). For survived tendons, 100,000 cycles were defined as the final fatigue stage because image quality deteriorated beyond this point. This cycle count exceeded the final fatigue life reached by all ruptured tendons while providing a consistent endpoint for comparison.

To evaluate local tissue deformation, regional longitudinal strain was calculated from the cumulative longitudinal displacement field. Longitudinal strain was estimated from the displacement gradient along the tendon axis using central finite differences between adjacent kernel columns under the small-strain assumption.^7,8^ For each interior column, the displacement difference between the adjacent columns on either side was divided by twice the column spacing. Strain values at the leftmost and rightmost columns were excluded from analysis because central finite differences could not be computed consistently at the boundaries. Longitudinal strain maps were generated at the peak loading frame for each fatigue stage. Maximum, average, and standard deviation of longitudinal strain were calculated within the strain map region to quantify strain magnitude and spatial heterogeneity. Principal strain directions were also determined to characterize local deformation orientation. Longitudinal strain maps were evaluated at eleven fatigue stages spanning approximately 1% to 100% fatigue life to characterize regional strain progression throughout fatigue loading. A second analysis was performed at eleven early fatigue stages ranging from 500 to 10,000 cycles, as these represented the most common number of cycles for all samples, to investigate whether regional strain behavior differed between groups before the onset of rupture.

One-way ANOVA with a Kruskal–Wallis correction was used to compare strain metrics between the survived and ruptured groups at the final fatigue stage. To evaluate the effects of fatigue progression, linear mixed-effects models (LME) were used to assess changes in regional strain metrics across fatigue life and during the early fatigue-loading period over the first 10,000 cycles. Group (survived or ruptured), fatigue progression (fatigue life or absolute cycle number), and their interaction were included as fixed effects, whereas tendon specimen was included as a random intercept. For the absolute-cycle analysis, cycle number was centered at 500 cycles and scaled per 1,000 cycles to facilitate interpretation of model coefficients. When significant interactions were detected, group-specific slopes were evaluated using linear contrast. Statistical significance was defined as p < 0.05. All statistical analyses were performed in MATLAB R2025b (MathWorks, Natick, MA, USA).

## 3. Results

### 3.1 Validation of Ultrasound-Derived Tendon Strain Measurements

Ultrasound-derived and the crosshead-derived tendon strains throughout the fatigue life were strongly correlated in both survived (R^2^ = 0.968 ± 0.017) and ruptured (R^2^ = 0.972 ± 0.014) groups (Fig. 2, Table 1). Similarly, the mean of root-mean-square error (RMSE) for the strain estimation remained low, below 0.050% in the survived group and below 0.070% in the ruptured group. Despite strong linear agreement, the relationship between ultrasound-derived and crosshead-derived tendon strain varied across specimens and fatigue stages. Group mean slopes were below unity in both survived and ruptured tendons, although several individual fatigue stages exhibited slopes greater than 1. Nevertheless, at the final fatigue stage, the ultrasound-derived tendon bulk strain was significantly greater in the ruptured group than in the survived group (Fig. 3).

**Table 1.** Summary of linear correlation metrics between ultrasound-derived tendon strain and crosshead-derived tendon strain across all cadaveric Achilles tendons.

| Sample ID | Age | Group | Slope (min – max) | $R^2$ (min – max) | Mean RMSE (% strain) |
| --- | --- | --- | --- | --- | --- |
| 045L | 63 | Survived | 0.36 - 1.10 | 0.982-0.994 | 0.026 |
| 127L | 59 | Survived | 0.45 - 1.01 | 0.962 -0.992 | 0.048 |
| 131R | 33 | Survived | 0.39 - 0.98 | 0.910- 0.986 | 0.050 |
| 153L | 55 | Survived | 0.16 - 0.69 | 0.915 - 0.987 | 0.036 |
| 125L | 75 | Failed | 0.59 - 1.36 | 0.966 - 0.993 | 0.044 |
| 125R | 75 | Failed | 0.20 - 1.74 | 0.900 - 0.994 | 0.047 |
| 127R | 59 | Failed | 1.05 - 2.33 | 0.963 - 0.994 | 0.063 |
| 140L | 58 | Failed | 0.11 - 0.90 | 0.855 - 0.993 | 0.041 |
| 153R | 55 | Failed | 0.45 - 1.04 | 0.922 - 0.991 | 0.070 |
| 176L | 78 | Failed | 0.73 - 1.00 | 0.973-0.994 | 0.042 |

| Group Summary | Mean Slope ± SD | Mean $R^2 \pm$ SD | Mean RMSE ± SD |
| --- | --- | --- | --- |
| **Survived** | 0.61 ± 0.09 | 0.968 ± 0.017 | 0.040 ± 0.012 |
| **Ruptured** | 0.95 ± 0.40 | 0.972 ± 0.014 | 0.051 ± 0.012 |

**Fig 2.**
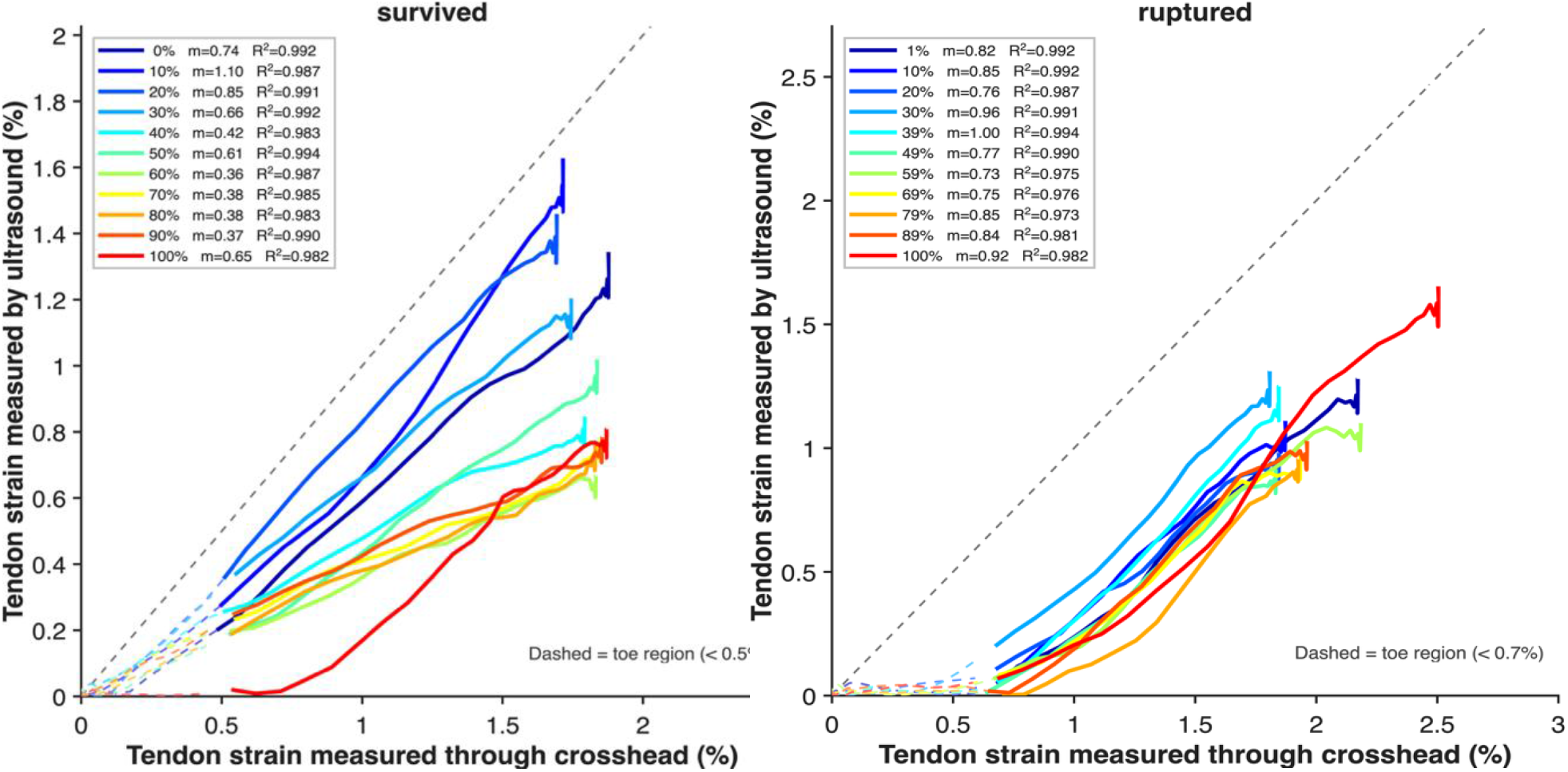
Representative correlation plots between ultrasound-derived tendon strain and actuator-derived tendon strain across fatigue life. (*Left*) Representative survived tendon. (*Right*) Representative failed tendon. Each colored curve represents a loading cycle at a different percentage of fatigue life. Solid lines indicate the linear region after removal of the toe region (dashed colored lines). The gray dashed diagonal represents the unity line (x = y). Linear fit slope (m) and coefficient of determination (R^2^) are shown for each fatigue stage.

**Fig 3.**
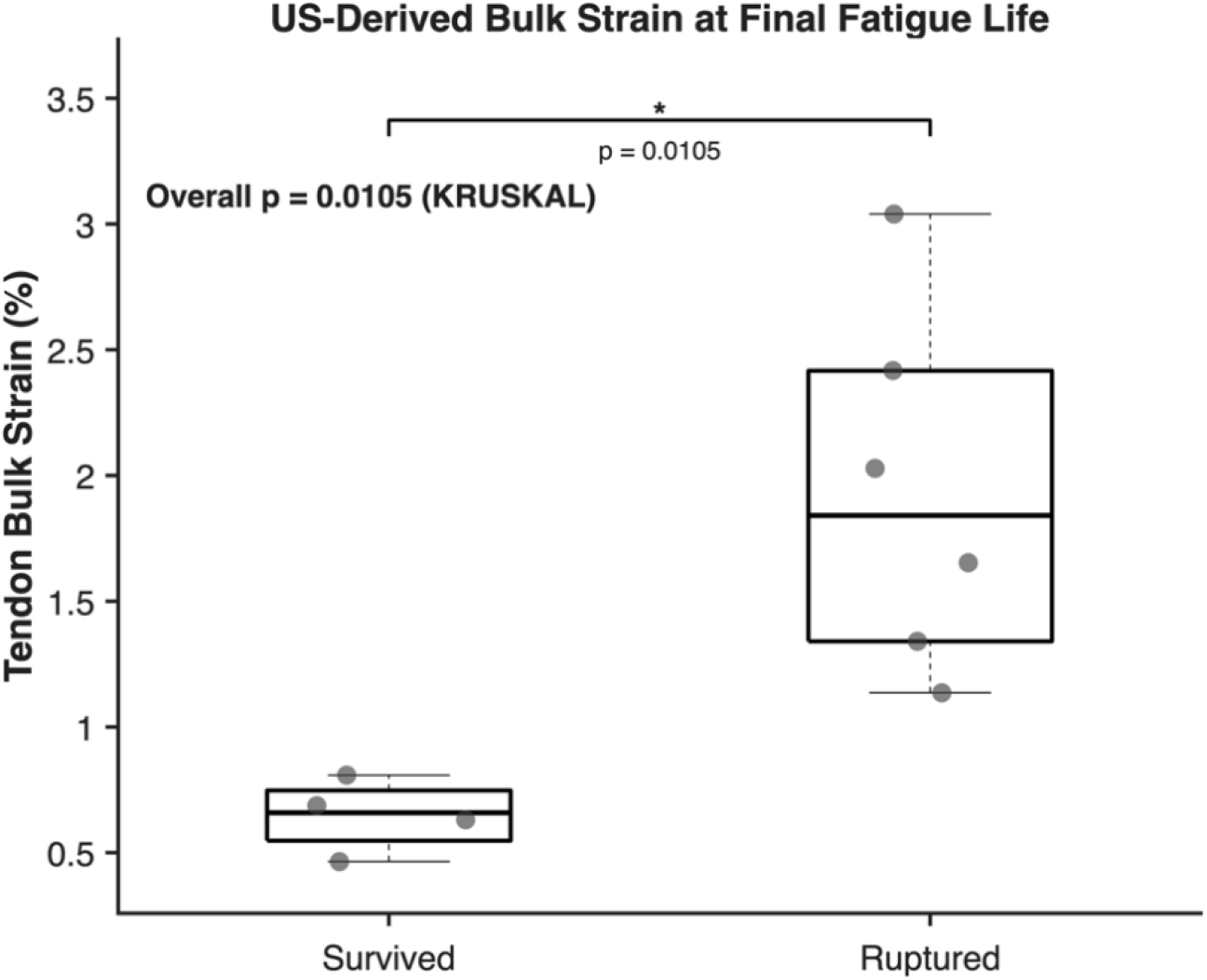
Tendon strain measured using ultrasound tracking at the final fatigue stage. The ruptured tendons exhibited significantly greater bulk strain compared with the survived tendons at the final fatigue cycle.

### 3.2 Regional Tendon Strains Throughout the Fatigue Life

Regional strain maps demonstrated evolving spatial strain behavior throughout fatigue life, with ruptured tendons exhibiting more localized regions of elevated longitudinal strain at later stages compared with survived tendons (Fig 4, Fig 5, and Table 2). Maximum longitudinal strain at peak stretch was significantly associated with fatigue life (p < 0.0001), and the significant group-by-fatigue-life interaction was observed (p = 0.003). Despite no overall group effect (p = 0.383), maximum strain evolved differently between groups; ruptured tendons exhibited persistently elevated strains, whereas survived tendons displayed decreasing strains over fatigue life (Fig 5A). Similarly, average longitudinal strain showed significant effects of both fatigue life (p = 0.0025) and group-by-fatigue-life interaction (p < 0.0001), with no overall group effect (p = 0.777). Average strain increased in the ruptured group over fatigue life but decreased in the survived tendons (Fig 5B). On the other hand, the standard deviation of longitudinal strain was significantly affected by the fatigue life (p < 0.0001), but not by the group-by-fatigue-life interaction (p = 0.146), or the overall group effect (p = 0.095), indicating reduced strain heterogeneity over fatigue life in both groups (Fig 5C).

**Table 2.** Linear mixed-effects model results for regional longitudinal strain metrics throughout normalized fatigue life.

| Outcome | Effect | Estimate | 95% CI | P-value |
| --- | --- | --- | --- | --- |
| Max Strain | Group | 1.255 | -1.588 to 4.099 | 0.383 |
|  | Fatigue Life | -0.046 | -0.063 to -0.030 | <0.001 |
|  | Group x Fatigue Life | 0.040 | 0.019 to 0.061 | <0.001 |
| Average Strain | Group | -0.064 | -0.508 to 0.380 | 0.777 |
|  | Fatigue Life | -0.004 | -0.007 to -0.001 | 0.003 |
|  | Group x Fatigue Life | 0.011 | 0.0080 to 0.015 | <0.001 |
| Strain SD | Group | 1.030 | -0.184 to 2.244 | 0.095 |
|  | Fatigue Life | -0.015 | -0.021 to -0.009 | <0.001 |
|  | Group x Fatigue Life | 0.006 | -0.002 to 0.014 | 0.146 |

**Fig 4.**
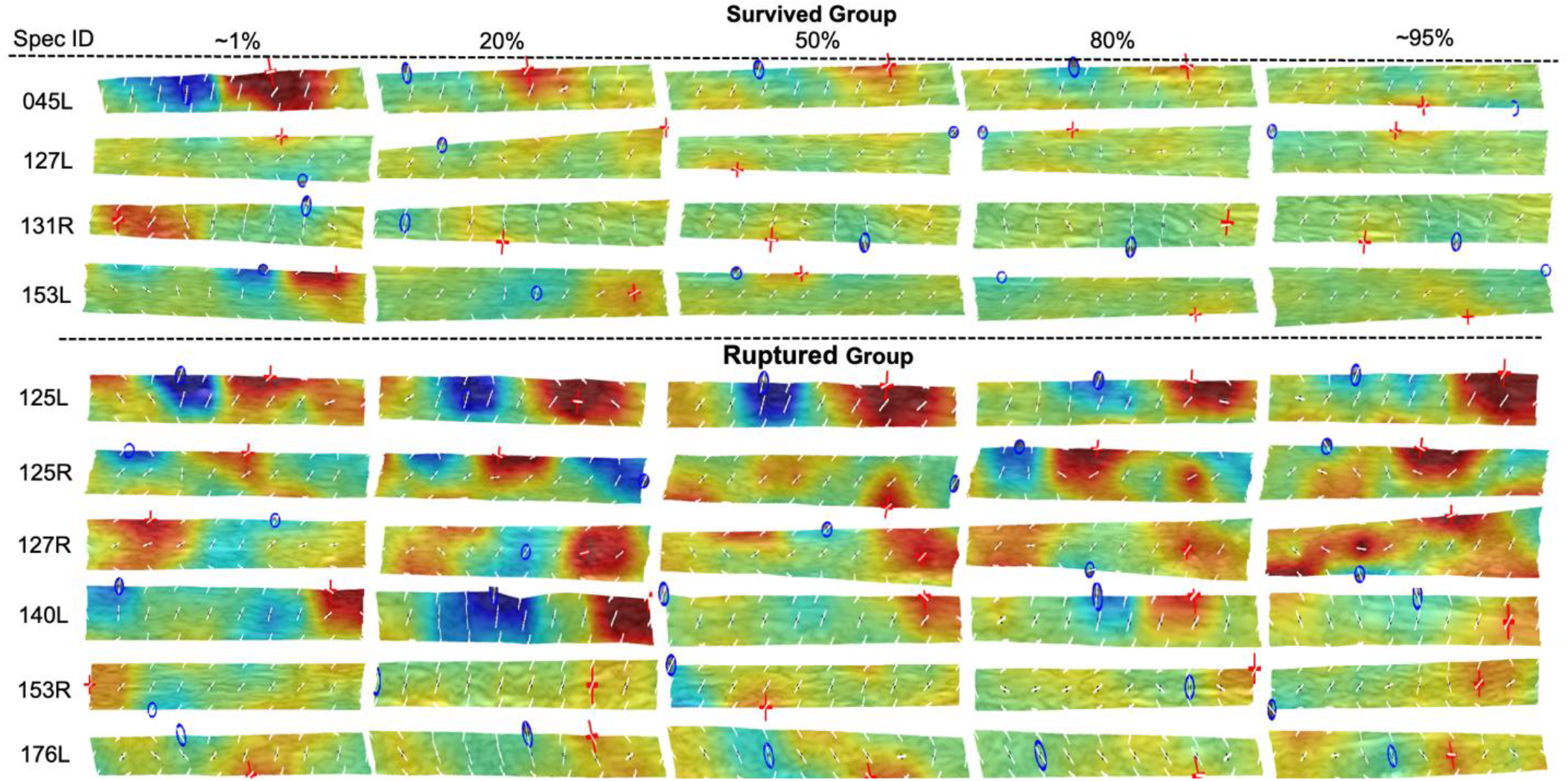
Longitudinal strain maps throughout fatigue life for all cadaveric Achilles tendons. Longitudinal strain maps are shown at 1%, 20%, 50%, 80%, and 99% of fatigue life for each tendon. Colors represent local longitudinal strain magnitude (% strain), with warmer colors indicating greater tensile strain, while white vectors indicate principal strain directions.

**Fig 5.**
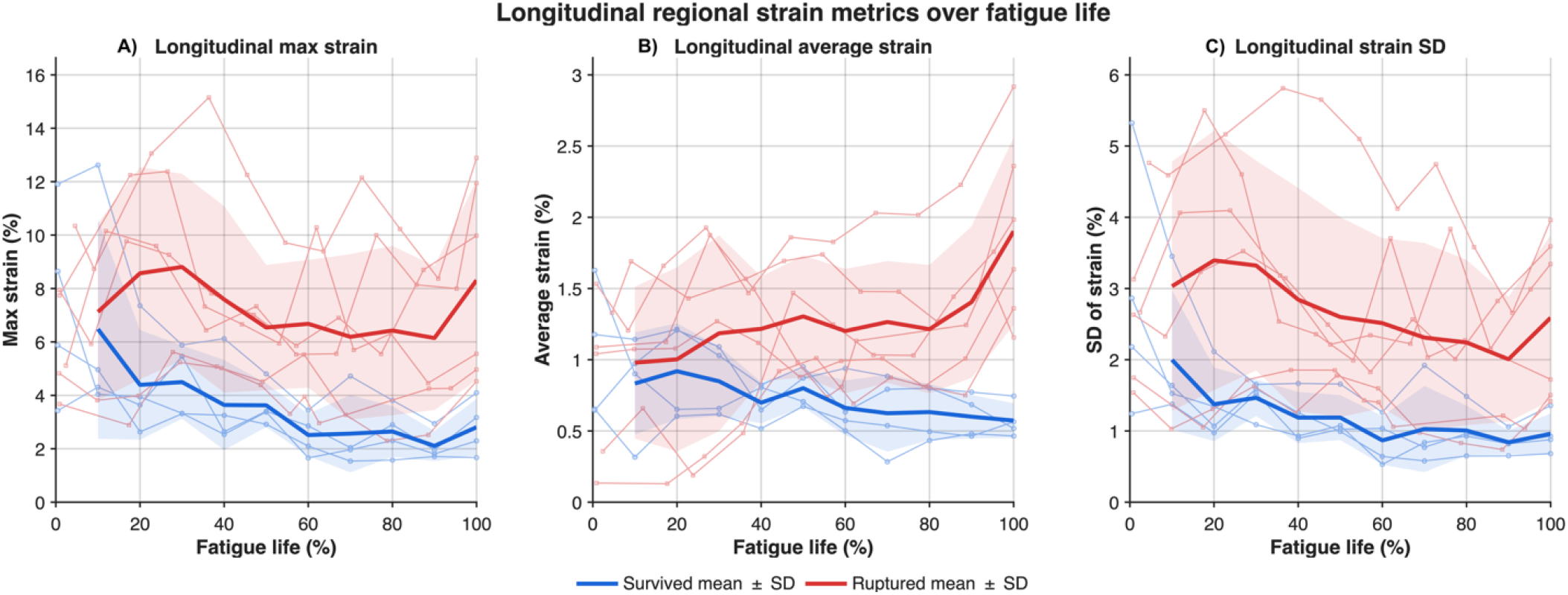
Longitudinal regional strain metrics over fatigue life for survived and ruptured tendons. Plots represent (A) maximum, (B) average, and (C) standard deviation of the longitudinal regional strain measured at the peak stretch frame across the fatigue life. Thick blue and red lines indicate the group means for the survived and ruptured tendons, respectively, while thin lines represent individual specimens. Shaded regions represent ±1 standard deviation of each group.

### 3.3 Regional Tendon Strain During Early Fatigue Life

During the first 10,000 loading cycles, there were no significant between-group differences in maximum, average, or standard deviation of longitudinal strain at the evaluated cycle counts (Table 3). However, average longitudinal strain exhibited a significant Group × Cycle interaction (p = 0.0061). Average strain decreased significantly with cycle number in survived tendons (β = −0.0268% per 1,000 cycles, p = 0.0215), whereas no significant change was observed in ruptured tendons (β = 0.0147% per 1,000 cycles, p = 0.1197) (Fig. 6B). Maximum longitudinal strain also decreased significantly with cycle number in survived tendons (β = −0.2215% per 1,000 cycles, p = 0.0222), but not in ruptured tendons (p = 0.5589). However, the Group × Cycle interaction was not significant (p = 0.1564) (Fig. 6A). Similarly, longitudinal strain standard deviation decreased significantly in survived tendons (β = −0.1070% per 1,000 cycles, p = 0.0029), but not in ruptured tendons (p = 0.2223), with no significant Group × Cycle interaction (p = 0.1157) (Fig. 6C).

**Table 3.** Linear mixed-effects model results for regional longitudinal strain metrics throughout 10,000 cycles.

| Outcome | Effect | Estimate | 95% CI | P-value |
| --- | --- | --- | --- | --- |
| Max Strain | Group | 1.420 | -2.834 to 5.674 | 0.510 |
|  | Survived Slope, %/1000 cycles | -0.222 | -0.411 to -0.032 | 0.022 |
|  | Failed Slope, %/1000 cycles | -0.046 | N/A | 0.559 |
|  | Group x Cycle Interaction | 0.176 | -0.068 to -0.420 | 0.156 |
| Average Strain | Group | 0.096 | -0.349 to 0.541 | 0.670 |
|  | Survived Slope, %/1000 cycles | -0.027 | -0.050 to -0.004 | 0.022 |
|  | Failed Slope, %/1000 cycles | 0.015 | N/A | 0.120 |
|  | Group x Cycle Interaction | 0.041 | 0.012 to 0.071 | 0.006 |
| Strain SD | Group | 0.728 | -1.076 to 2.532 | 0.426 |
|  | Survived Slope, %/1000 cycles | -0.107 | -0.177 to -0.037 | 0.003 |
|  | Failed Slope, %/1000 cycles | -0.035 | N/A | 0.222 |
|  | Group x Cycle Interaction | 0.072 | -0.018 to 0.162 | 0.116 |

**Fig 6.**
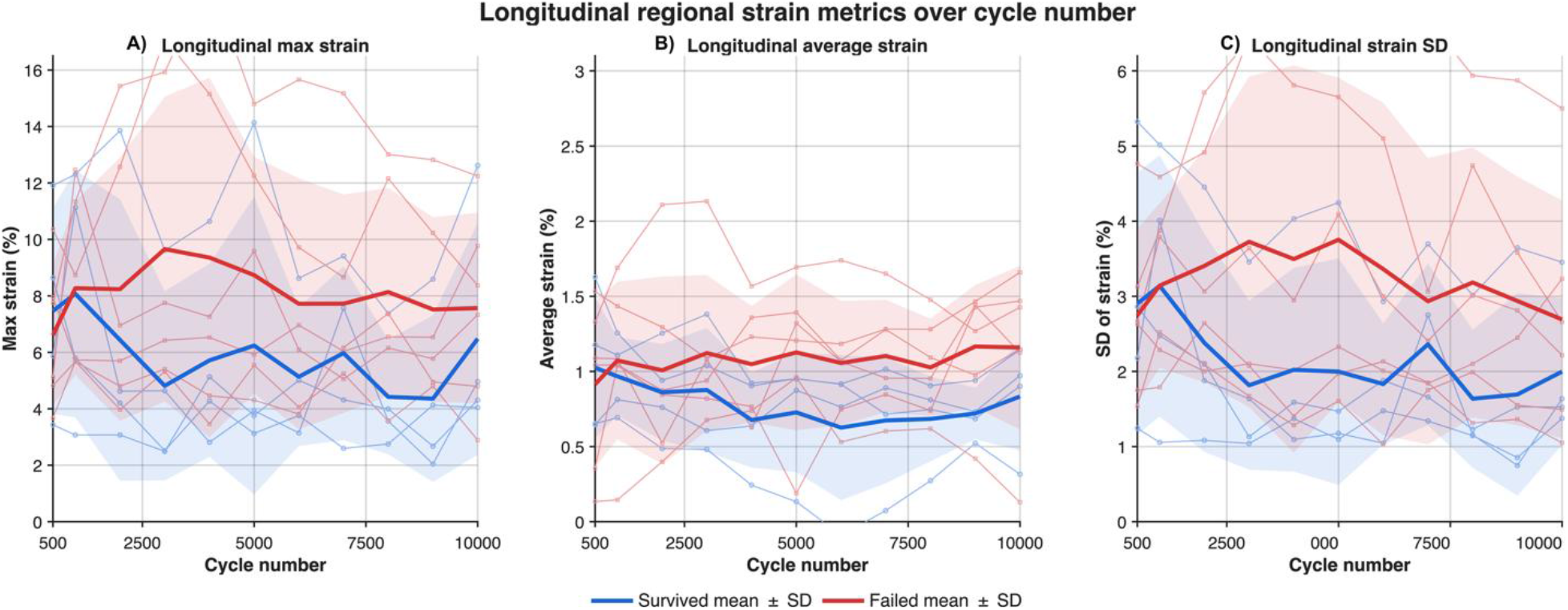
Longitudinal regional strain metrics over fatigue life for survived and ruptured tendons. Plots represent maximum, average, and standard deviation of the longitudinal regional strain measured at the peak stretch frame across the fatigue life. Thick blue and red lines indicate the group means for the survived and ruptured tendons, respectively, while thin lines represent individual specimens. Shaded regions represent ±1 standard deviation of each group.

## 4. Discussion

This study developed and validated a robust automated tracking algorithm using Kanade-Lucas-Tomasi algorithm to quantify tendon displacement and assess regional tendon strain throughout progressive fatigue loading. The ultrasound-based tracking algorithm demonstrated high tracking fidelity when ultrasound-derived bulk tendon strain was compared with crosshead-derived strain across the fatigue life in both survived and ruptured tendons. Furthermore, our regional strain analysis revealed that maximum and average longitudinal strains evolved differently between groups, with progressive divergence occurring as fatigue loading advanced. In contrast, no early differences in regional strains were detected between groups. Therefore, our hypothesis that regional strain progressively increases in ruptured tendons was partially supported. Collectively, these findings validate our automated tracking algorithm and provide insight into the evolution of regional strain behaviors during fatigue loading.

We validated our ultrasound-based tracking algorithm by comparing ultrasound-derived bulk tendon strain with crosshead-derived tendon strain from our previous study.^33^ The strong correspondence between the two measurements in both survived and ruptured tendons supports the validity of the KLT-based approach for tracking tendon deformation throughout fatigue loading. Previous B-mode ultrasound studies have demonstrated accurate tendon displacement and strain measurements, with Korstanje et al. reporting mean relative displacement errors of approximately 1.0%–1.6% across cadaveric and in vivo experiments and Okotie et al. reporting a near-unity slope between ultrasound texture and optical marker-derived tendon strains.^23,30^ This study extends these validation efforts by evaluating tendon strain throughout fatigue loading and across different tendon outcomes. The consistently high correlation values across fatigue stages indicate that the algorithm reliably captured changes in tendon strain across loading, while low RMSE values indicate small absolute discrepancies between these measurements (Table 1). Although the group mean slopes were below unity in both groups, individual slopes spanned values below and above unity across specimens and fatigue stages. The tendency toward slopes below unity is consistent with previous B-mode speckle-tracking studies that reported modest underestimation of tendon motion and strain.^4,7,8^ The tendency toward below-unity slopes was expected, in part, because of testing-system compliance, as crosshead displacement captured deformation of the grips, cemented fixtures, and other components of the mechanical testing system in addition to elongation of the tendon specimen. In contrast, slopes above unity may reflect strain concentration within the narrow midsubstance of the dog-bone specimens, where ultrasound measurements were obtained, relative to the more global strain measured across the entire specimen by the crosshead.^7,8^ Therefore, this variability in individual slope may be attributed to the combined effects of typical B-mode motion estimation, the specimen geometry, and the differences between localized ultrasound and whole-specimen crosshead measurements. Importantly, the algorithm detected greater final-stage tendon bulk strain in the ruptured group than the survived group, consistent with our crosshead-based findings.^33^ The consistency suggests that the KLT-based approach still preserved biologically meaning between-group differences despite variability in absolute strain magnitude. Taken together, these findings support the use of KLT-based ultrasound tracking to characterize tendon deformation and regional strain progression during fatigue loading.

Regional longitudinal strains evolved differently in tendons that ultimately ruptured compared with those tendons that survived fatigue loading. Maximum longitudinal strain and average longitudinal strain progressively diverged between groups, with the ruptured group exhibiting greater strain at the later fatigue stages. These diverging trajectories were supported by significant Group x FatigueLife interactions. Previous ultrasound-based studies have demonstrated ability to characterize nonuniform intratendinous displacement and regional strain. Borgaerts et al. evaluated regional strain within the superficial, middle, and deep layers of the Achilles tendons and found greater displacement in the deep layer and the greater regional strain in the superficial layer during passive elongation.^5^ Extending this layer-based analysis, the present study implemented a denser grid of tracked kernels to generate spatially regional strain maps that allow characterizing strain magnitude and heterogeneity throughout fatigue loading. In ruptured tendons, maximum regional strain values in the later fatigue stage reached above 8%, which lie within reported failure range for human Achilles tendons.^15^ However, this magnitude should not be interpreted as a threshold for macroscopic failure since in vivo ultrasound studies have reported 5% - 8% free Achilles tendon strain during maximal voluntary contraction.^27–29^ Therefore, the local strain maxima in ruptured tendons may represent concentrated deformation resulting from advanced mechanical deterioration. Previous studies have demonstrated that repetitive loading leads to progressive subrupture fatigue damage, starting from kinked fiber deformation to increasing fiber dissociation, discontinuities, and localized rupture.^20^ These microstructural changes reduce the number of intact fibers available to support the applied load resulting in the diminishing tendon stiffness and load-bearing capacity.^19,39^ Consistent with this mechanism, Achilles tendons subjected to cyclic loading have demonstrated progressive reductions in modulus as fatigue damage accumulates. Therefore, microstructural damage and mechanical degradation may explain both increasing maximum and average regional strains and the greater final stage bulk strain in the ruptured tendons. In terms of strain heterogeneity, standard deviation of the longitudinal strain decreased across fatigue life. However, neither the group effect nor the Group x FatigueLife interaction was the significant, indicating that the magnitude and trajectory of this decrease were similar between these groups. These findings suggest that tendon rupture risk is more strongly reflected by changes in maximum and average regional strain than by the spatial strain heterogeneity. To determine whether these differences were also evident before fatigue damage accumulated, regional strain pattern was investigated during the first 10,000 cycles. Although there were no differences between groups at individual cycles, average longitudinal strain exhibited divergent trajectories between groups, decreasing in survived tendons while remaining relatively unchanged in ruptured tendons. Maximum longitudinal strain and strain heterogeneity also decreased in survived tendons, despite no statistical difference in trajectories from those of ruptured tendons. This early decline in average strain in survived tendons may reflect mechanical conditioning and load redistribution during repetitive loading. In contrast, the absence of a similar downward trend in ruptured tendons may suggest an altered early biomechanical response to cyclic loading before progressively stark differences in the later fatigue stages. Collectively, these findings partially supported our hypotheses, with maximum and average strains diverging between groups across fatigue life and average strain exhibiting different trajectories during the first 10,000 cycles, whereas strain heterogeneity did not significantly differ between groups.

Ultrasound is widely used to evaluate tendon morphology in vivo, while ultrasound-based measurements of Achilles tendon mechanical behavior have increasingly been investigated for tendon health assessment and monitoring. For instance, Konggaard et al. demonstrated reproducible measurements of Achilles tendon deformation and stiffness across two testing sessions performed one week apart.^22^ Characterization of Achilles tendon mechanical properties may provide information about tendon pathology beyond structural morphology alone. Previous studies have reported altered tendon strain in active individuals with Achilles tendinopathy and during passive ankle dorsiflexion in individuals with insertional tendinopathy.^9,10^ Furthermore, Arya and Kulig similarly reported greater compliance in tendinopathic Achilles tendons using ultrasound-based tracking approach.^2^ At a more regional level, ultrasound speckle tracking has been used to characterize the degree of differential displacements within Achilles tendon. Couppe et al. reported decreased differential displacement in tendinopathic tendons compared with the asymptomatic contralateral side, while Froberg et al. observed similarly reduced nonuniform displacement following surgical repair of Achilles tendon rupture.^12,18^ In the present study, the KLT-based tracking algorithm generated spatially resolved regional strain fields using a dense grid of tracked kernels, enabling quantification of the maximum, average, and standard deviation of regional strain. The algorithm also revealed different early trajectories in average regional strain between tendons that ultimately ruptured and those that survived. These findings suggest potential utility for identifying tendons with altered mechanical behavior before pronounced differences in strain magnitude develop. Although the present study was not designed to establish rupture-prediction thresholds, longitudinal changes in regional strain may warrant further investigation as candidate biomechanical markers of future tendon failure. Therefore, these spatially resolved biomechanical metrics may complement existing ultrasound assessments by supporting longitudinal monitoring of local tendon mechanics, identifying regions of concentrated deformation, and evaluating mechanical changes during rehabilitation.

This study has several limitations related to the ultrasound imaging approach, the ex vivo fatigue model, specimen preparation, and the relatively small and biologically heterogeneous sample. First, although longitudinal, transverse, and shear strain components could be estimated within the imaging plane, sagittal-plane ultrasound provided only a two-dimensional representation of tendon mechanics and could not capture out-of-plane displacement or the complete three-dimensional strain state. In addition, the restricted ultrasound field of view limited the analysis to the tendon midsubstance and prevented characterization of deformation outside the imaged region. The algorithm also applied temporal median and spatial filtering to improve displacement-tracking robustness; however, these procedures may have attenuated local displacement gradients and consequently underestimated peak regional strain and strain heterogeneity. Second, the controlled uniaxial loading used in the ex vivo model could not reproduce muscle-driven Achilles tendon mechanics in vivo. In vivo tendon deformation is influenced by the differential contributions of the medial gastrocnemius, lateral gastrocnemius, and soleus muscles, as well as by joint posture, surrounding tissues, and tendon constraints, resulting in nonuniform intratendinous displacement.^11^ Third, freezing, storage, and dissection into a dog-bone geometry may have altered tissue behavior before mechanical testing. Although the narrowed geometry promoted deformation and localized failure within the imaged midsubstance, the resulting absolute strain magnitudes may not directly represent intact Achilles tendons in vivo. Variations in gauge width and thickness, irregular cut edges, specimen alignment, and clamping may also have introduced localized stress concentrations and contributed to the observed strain heterogeneity. Because the ultrasound field of view did not extend to the clamps or the transitions into the narrowed gauge region, these boundary effects could not be directly evaluated. Fourth, ruptured tendons failed at or below approximately 60,000 cycles at most, whereas survived tendons completed 150,000 cycles. We selected 100,000 cycles as the normalization end point for survived tendons because ultrasound image quality was consistently adequate through this stage, with fewer artifacts than later cycles, while still representing a loading duration beyond failure range of the ruptured tendons. Our prior analyses that quantified changes in image brightness found stable measurements between 100,000 to 150,000 cycles suggesting that tendon damage was not progressing in these survived tendons. Consequently, because percentage fatigue life was normalized to different endpoints across specimens, equivalent percentages did not necessarily correspond to the same absolute number of loading cycles. The complementary early-cycle analysis partially addressed this limitation by comparing tendons at common absolute cycle numbers. Finally, only ten specimens were available, and variability in donor age, sex, baseline tissue quality, and fatigue life may have reduced statistical power and limited the generalizability of the findings. Future work should incorporate wider-field or three-dimensional ultrasound imaging, more standardized specimen preparation, and a larger and more balanced cohort, followed by validation under in vivo loading conditions.

This study introduced an automated ultrasound-based tracking approach that quantified tendon deformation with high tracking fidelity and generated spatially resolved regional strain maps throughout the fatigue loading. The maximum and average longitudinal stains progressively diverged between ruptured and survived tendons across fatigue stages, while average longitudinal strain also exhibited different trajectories between groups during the first 10,000 loading cycles. These finding suggest that magnitude and evolution of regional strain may provide a useful biomarker for progressive mechanical deterioration. Altogether, this framework establishes a foundation for future studies in assessment of regional tendon mechanics during exercise, injury, and rehabilitation.

## Acknowledgments

We thank Mr. Todd Hullfish for help with data organization.

## References

1. Arndt A, Bengtsson AS, Peolsson M, Thorstensson A, Movin T. Non-uniform displacement within the Achilles tendon during passive ankle joint motion. Knee Surg Sports Traumatol Arthrosc. 2012;20(9):1868–1874.

2. Arya S, Kulig K. Tendinopathy alters mechanical and material properties of the Achilles tendon. J Appl Physiol. 2010;108(3):670–675.

3. Astrom M, Gentz CF, Nilsson P, Rausing A, Sjoberg S, Westlin N. Imaging in chronic achilles tendinopathy: a comparison of ultrasonography, magnetic resonance imaging and surgical findings in 27 histologically verified cases. Skeletal Radiology. Published online 1996.

4. Beyer R, Agergaard AS, Magnusson SP, Svensson RB. Speckle tracking in healthy and surgically repaired human Achilles tendons at different knee angles—A validation using implanted tantalum beads. TRANSLATIONAL SPORTS MEDICINE. 2018;1(2):79–88.

5. Bogaerts S, De Brito Carvalho C, Scheys L, et al. Evaluation of tissue displacement and regional strain in the Achilles tendon using quantitative high-frequency ultrasound. PLoS One. 2017;12(7):e0181364.

6. Bullock M, Pierson Z. Achilles Tendon Rupture. Clinics in Podiatric Medicine and Surgery. 2024;41(3):535–549.

7. a. Chernak L, Thelen DG. Tendon motion and strain patterns evaluated with two-dimensional ultrasound elastography. Journal of Biomechanics. 2012;45(15):2618–2623.

8. Chernak Slane L, Thelen DG. The use of 2D ultrasound elastography for measuring tendon motion and strain. J Biomech. 2014;47(3):750–754.

9. Child S, Bryant AL, Clark RA, Crossley KM. Mechanical properties of the achilles tendon aponeurosis are altered in athletes with achilles tendinopathy. Am J Sports Med. 2010;38(9):1885–1893.

10. Chimenti RL, Bucklin M, Kelly M, et al. Insertional Achilles tendinopathy associated with altered transverse compressive and axial tensile strain during ankle dorsiflexion. J Orthop Res. 2017;35(4):910–915.

11. Clark WH, Franz JR. Do triceps surae muscle dynamics govern non-uniform Achilles tendon deformations? PeerJ. 2018;6(e5182):e5182.

12. Couppé C, Svensson RB, Josefsen CO, Kjeldgaard E, Magnusson SP. Ultrasound speckle tracking of Achilles tendon in individuals with unilateral tendinopathy: a pilot study. Eur J Appl Physiol. 2020;120(3):579–589.

13. Crouzier M, Baudry S, Vanwanseele B. Achilles Subtendons Stiffness Differ in People with and without Achilles Tendinopathy. Medicine & Science in Sports & Exercise. 2025;57(8):1636–1645.

14. Docking SI, Rio E, Cook J, Carey D, Fortington L. Quantification of Achilles and patellar tendon structure on imaging does not enhance ability to predict self-reported symptoms beyond grey-scale ultrasound and previous history. Journal of Science and Medicine in Sport. 2019;22(2):145–150.

15. Doral MN, Alam M, Bozkurt M, et al. Functional anatomy of the Achilles tendon. Knee Surgery, Sports Traumatology, Arthroscopy. 2010;18(5):638–643.

16. Drazan JF, Hullfish TJ, Baxter JR. An automatic fascicle tracking algorithm quantifying gastrocnemius architecture during maximal effort contractions. PeerJ. 2019;7(e7120):e7120.

17. Franz JR, Slane LC, Rasske K, Thelen DG. Non-uniform in vivo deformations of the human Achilles tendon during walking. Gait Posture. 2015;41(1):192–197.

18. Fröberg Å, Cissé AS, Larsson M, et al. Altered patterns of displacement within the Achilles tendon following surgical repair. Knee Surg Sports Traumatol Arthrosc. 2017;25(6):1857–1865.

19. Fung DT, Wang VM, Andarawis-Puri N, et al. Early response to tendon fatigue damage accumulation in a novel in vivo model. J Biomech. 2010;43(2):274–279.

20. Fung DT, Wang VM, Laudier DM, et al. Subrupture tendon fatigue damage. J Orthop Res. 2009;27(2):264–273.

21. Jolliffe IT, Cadima J. Principal component analysis: a review and recent developments. Philos Trans A Math Phys Eng Sci. 2016;374(2065):20150202.

22. Kongsgaard M, Nielsen CH, Hegnsvad S, Aagaard P, Magnusson SP. Mechanical properties of the human Achilles tendon, in vivo. Clin Biomech (Bristol, Avon). 2011;26(7):772–777.

23. Korstanje JWH, Selles RW, Stam HJ, Hovius SER, Bosch JG. Development and validation of ultrasound speckle tracking to quantify tendon displacement. J Biomech. 2010;43(7):1373–1379.

24. Lantto I, Heikkinen J, Flinkkila T, et al. A prospective randomized trial comparing surgical and nonsurgical treatments of acute Achilles tendon ruptures. Am J Sports Med. 2016;44(9):2406–2414.

25. Lucas BD, Kanade T. An Iterative Image Registration Technique with an Application to Stereo Vision. In: IJCAI’81: 7th International Joint Conference on Artificial Intelligence. Vol 2. ; 1981:674–679.

26. Maffulli N, Longo UG, Maffulli GD, Khanna A, Denaro V. Achilles Tendon Ruptures in Elite Athletes. Foot Ankle Int. 2011;32(1):9–15.

27. Maganaris CN, Narici MV, Maffulli N. Biomechanics of the Achilles tendon. Disabil Rehabil. 2008;30(20-22):1542–1547.

28. Maganaris CN, Paul JP. Tensile properties of the in vivo human gastrocnemius tendon. J Biomech. 2002;35(12):1639–1646.

29. Magnusson SP, Hansen P, Aagaard P, et al. Differential strain patterns of the human gastrocnemius aponeurosis and free tendon, in vivo. Acta Physiol Scand. 2003;177(2):185–195.

30. Okotie G, Duenwald-Kuehl S, Kobayashi H, Wu MJ, Vanderby R. Tendon Strain Measurements With Dynamic Ultrasound Images: Evaluation of Digital Image Correlation. J Biomech Eng. 2012;134(2):24504–NaN.

31. Peng WC, Chao YH, Fu ASN, et al. Muscular morphomechanical characteristics after an Achilles repair. Foot Ankle Int. 2019;40(5):568–577.

32. van Schie HTM, Vos RJ de, Jonge S de, et al. Ultrasonographic tissue characterisation of human Achilles tendons: quantification of tendon structure through a novel non-invasive approach. Published online December 1, 2010.

33. Schmidt EC, Hullfish TJ, O’Connor KM, Hast MW, Baxter JR. Ultrasound echogenicity is associated with fatigue-induced failure in a cadaveric Achilles tendon model. J Biomech. 2020;105(109784):109784.

34. Shi J, Tomasi. Good features to track. In: 1994 Proceedings of IEEE Conference on Computer Vision and Pattern Recognition. ; 1994:593–600.

35. Slane LC, Thelen DG. Non-uniform displacements within the Achilles tendon observed during passive and eccentric loading. J Biomech. 2014;47(12):2831–2835.

36. Svensson RB, Couppé C, Agergaard AS, et al. Persistent functional loss following ruptured Achilles tendon is associated with reduced gastrocnemius muscle fascicle length, elongated gastrocnemius and soleus tendon, and reduced muscle cross-sectional area. TRANSLATIONAL SPORTS MEDICINE. 2019;2(6):316–324.

37. Szczesny SE, Aeppli C, David A, Mauck RL. Fatigue Loading of Tendon Results in Collagen Kinking and Denaturation but Does Not Change Local Tissue Mechanics. J Biomech. 2018;71:251–256.

38. Tomasi, C. and Kanade, T. (1991) Detection and Tracking of Point Features Technical Report CMU-CS-91-132. Image Rochester, NY. 1991, 91(April), 1–22. - References - Scientific Research Publishing. n.d.

39. Wren TAL, Lindsey DP, Beaupré GS, Carter DR. Effects of creep and cyclic loading on the mechanical properties and failure of human Achilles tendons. Ann Biomed Eng. 2003;31(6):710–717.

